# Homologous recombination substantially delays sequence but not gene content divergence of prokaryotic populations

**DOI:** 10.1101/518852

**Authors:** Jaime Iranzo, Yuri I. Wolf, Eugene V. Koonin, Itamar Sela

## Abstract

Evolution of bacterial and archaeal genomes is a highly dynamic process that involves extensive gain and loss of genes. Therefore, phylogenetic trees of prokaryotes can be constructed both by the traditional sequence-based methods (gene trees) and by comparison of gene compositions (genome trees). Comparing the branch lengths in gene and genome trees with identical topologies for 34 clusters of closely related bacterial and archaeal genomes, we found that the terminal branches of gene trees were systematically compressed compared to those of genome trees. Thus, sequence evolution seems to be significantly delayed with respect to genome evolution by gene gain and loss. The extent of this delay widely differs among bacterial and archaeal lineages. We develop and explore mathematical models demonstrating that the delay of sequence divergence can be explained by sequence homogenization that is caused by homologous recombination. Once evolving genomes become isolated by barriers that impede homologous recombination, gene and genome evolution processes settle into parallel trajectories, and genomes diverge, resulting in speciation. This model of prokaryotic genome evolution gives a mechanistic explanation of our previous finding that archaeal genomes contain a class of genes that turn over rapidly, before significant sequence divergence occurs, and provides a framework for correcting phylogenetic trees, to make them consistent with the dynamics of gene turnover.

## Introduction

Evolution of bacterial and archaeal genomes is a highly dynamic process that involves extensive gain and loss of genes, with turnover rates comparable to if not exceeding the rate of nucleotide substitution^1–3^. Gene gain and loss occur through insertion and deletion of genome segments of variable size, including large genomic islands, via mechanisms of non-homologous recombination, often involving mobile genetic elements^4,5^. The gene gain and loss events can be used to generate “gene content trees” that reflect the evolution of microbial pangenomes and complement traditional phylogenetic trees constructed from sequence alignments of highly conserved marker genes. From such trees, a gene turnover clock can be defined. The gene turnover clock ticks at a rate that does not necessarily correlate with the rate of the traditional, sequence-based molecular clock.

Evolution of prokaryotic populations is strongly affected by homologous recombination, which is regarded as a major contributor to maintaining genetic cohesion by preventing sequence divergence via gene conversion^6,7^. However, because barriers to recombination do not necessarily affect the whole genome, bacterial strains can diverge at some loci while remaining cohesive at others ^8^. The rate of population divergence and, eventually, speciation thus depends on the dynamics of recombination barrier emergence across genomes.

Efficient homologous recombination between two genomes requires the presence of (nearly) identical nucleotide sequences flanking the exchanged genomic regions. The minimum length of these flanks depends on the species, with typical values around 25-100 nucleotides^9,10^. As genomes diverge, the probability to find fully conserved flanking sequences decreases, and so does the efficiency of recombination^11,12^. The existence of genetic barriers to homologous recombination was initially observed in experimental studies which have shown that sequence divergence of over 5% can prevent most recombination events in some bacteria^13–15^. Subsequently, comparative genomic analyses have confirmed that barriers to homologous recombination are widespread and can be used to define biological species in bacteria and archaea^16–18^. Mechanistic modeling of the molecular processes involved in homologous recombination has shown that barriers to recombination can build up spontaneously if the balance of mutation and recombination favors the sequence variability in the population^19–21^. Barriers to recombination can also arise after the acquisition of new genes^8,22^, especially if the newly acquired genes are involved in niche specialization^23,24^. In this case, the barriers seem to result, in part, from selection against gene conversion events that would lead to the loss of the recently acquired, beneficial genes.

Here, we investigate how the fraction of genes shared by closely related bacterial and archaeal genomes decays with the phylogenetic distance and show that molecular and gene turnover clocks are incongruent at short evolutionary times. To elucidate the origin of this discrepancy, we develop a mathematical model of genome evolution that describes the dynamics of sequence divergence in the presence of gene conversion. The model predicts the existence of a recombination-driven delay in the molecular clock, the magnitude of which corresponds to the time required for barriers to recombination to spread across the genome. By fitting the model to genomic data, we obtain estimates of such recombination-driven delays in 34 groups of closely related bacteria and archaea, and show that the incongruence between the molecular and gene turnover clocks disappears if the former is corrected to account for the estimated delay. Finally, we investigate the tempo and the factors that contribute to the establishment of barriers to recombination in the populations of diverse bacteria and archaea.

## Results

### Discrepancy between the molecular clock and the gene turnover clock

The evolution of gene content in closely related genomes has been recently investigated by mathematical modeling and comparative genomics^25–28^. These analyses have shown that the fraction of genes shared by a pair of genomes decays exponentially with time as the genomes diverge, and that genes can be classified in two categories based on their turnover rates^26^. Here we extend these approaches to sets of 3 and more genomes. For illustration, let us consider a simple scenario in which all genes have the same turnover rate *λ*. The fraction of genes shared by a pair of genomes is then

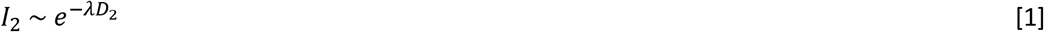

where *D*_2_ is the total evolutionary tree distance, which in the case of 2 genomes is equal to twice the distance from the last common ancestor. We show in the Methods that a similar formula describes the divergence in gene content for groups of 3 or more genomes. Specifically, the fraction of genes shared by *k* genomes decays as

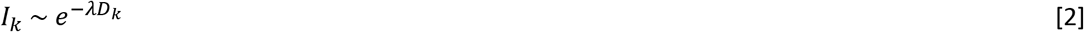

where *D_k_* is the total evolutionary distance spanned by those *k* genomes. Given a phylogenetic tree, the total distance *D_k_* is the sum of branch lengths of the subtree that includes the *k* genomes. The most notable aspect of this result is that the dynamics of gene content divergence is independent of the number of genomes considered (*k*). As a result, plots of the fraction of shared genes (*I_k_*) as a function of the total evolutionary distance (*D_k_*) for different sample sizes collapse into a single curve (Fig. 1a). This property remains valid under very general models of gene turnover, including the case where the rates of gene gain and loss differ across gene families (see Methods), under the condition that the tree distance is proportional to the rate of the gene turnover clock.

**Figure 1.**
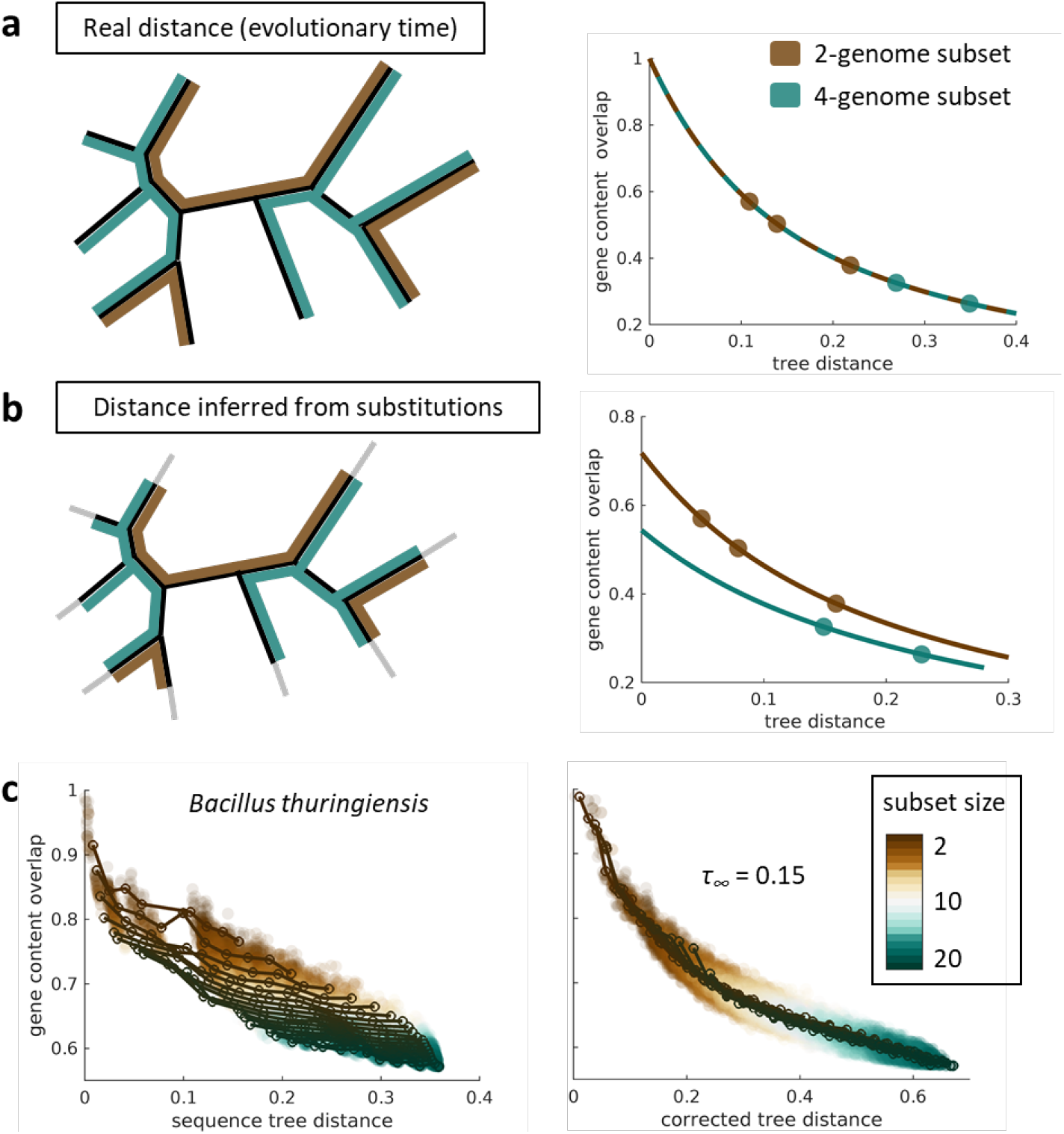
Homologous recombination leads to compression of terminal branches in sequence-based phylogenetic trees that can be detected through the analysis of gene content decay curves. a) If tree distances are proportional to the true evolutionary time, the fraction of genes shared by a subset of genomes will decay with the total length of the subtree, and the decay curves will be the same regardless of the number of genomes in the subset. For illustration purposes, three subsets of 2 genomes are highlighted in brown, and two subsets of 4 genomes are highlighted in green. b) Homologous recombination between pairs of closely related genomes erases recent sequence divergence which results in an underestimation of the evolutionary times associated with terminal tree branches. Such underestimation leads to gene content decay curves that depend on the number of genomes included in the subset. Accordingly, the decay curve of subsets of 4 genomes is different from the decay curve of subsets of 2 genomes. c) The gene content decay curves of the *Bacillus thuringiensis/ cereus/ anthracis* group are compatible with a scenario of recombination-driven shortening of the terminal tree branches (left plot, based on the tree from Fig. 2a). On the right, if the recombination model is used to correct for unobserved variation (fit in Fig. 2c, left panel), overlapping decay curves are obtained.

To test this theoretical prediction, we analyzed the profiles of gene sharing in 34 groups of closely related genomes from Bacteria and Archaea. As a proxy for the evolutionary time, we used the branch lengths of high-resolution sequence similarity phylogenetic trees built from concatenated alignments of single-copy core genes. Then, we sampled subsets of genomes and represented the fraction of shared genes as a function of the total tree distance. At odds with the theoretical expectation, we found that gene-sharing decay curves depend on the number of sampled genomes: as more genomes are added, the curves shift down and the fraction of shared genes becomes smaller than expected (left panel on Fig. 1c shows a representative case; see also Supplementary Fig. S1). As illustrated by Fig. 1b, such a non-overlapping pattern could be easily explained if the lengths of the terminal branches in the phylogenetic trees were systematically underestimated. In more general terms, the curves in Fig. 1c involve two different clocks: the molecular clock, that is used to infer the branch lengths in the phylogenetic tree; and the gene turnover clock, that governs the stochastic process of gene loss and, with it, the decay in the fraction of shared genes. Thus, the absence of overlap among gene sharing curves reflects a non-linear relationship between these two clocks, or more specifically, a delay in the molecular clock relative to the gene turnover clock.

To further explore the differences between the molecular clock and the gene turnover clock, we generated an alternative set of phylogenetic trees by rescaling branch lengths so that they are proportional to the number of gene gains and losses occurred during the evolution of a lineage (Fig. 2a,b; see Methods for details). Such “gene content trees” are topologically congruent with the gene turnover clock, as long as the genomes are close enough to assume that gene turnover rates have remained approximately constant since the last common ancestor (this approach does not require, however, that gain and loss rates are homogeneous across genes). The non-linear relationship between the molecular clock and the gene turnover clock becomes evident when comparing pairwise distances among leaves in both sets of trees (Fig. 2c). Sequence divergence only starts building up after a transient period during which genes are gained and lost, although the duration of this transient phase varies across taxa.

**Figure 2.**
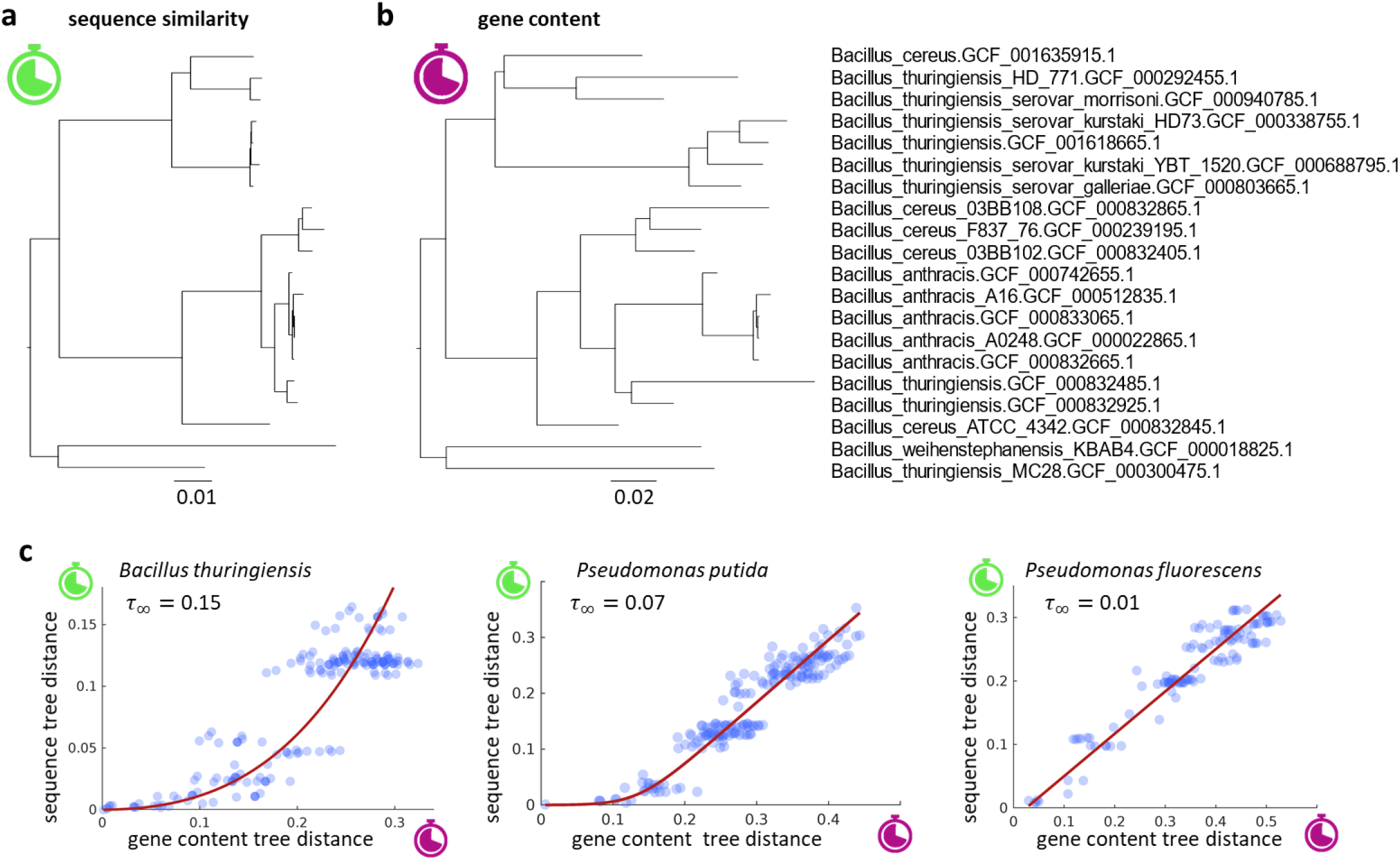
Lack of collinearity between sequence and gene content distances supports a recombination-drive delay in the molecular clock. a) Phylogenetic tree of the *Bacillus thuringiensis/cereus/anthracis* group based on the concatenated alignment of genes shared by all members of the group. b) The same tree, with branch lengths proportional to the number of gene gain and loss events estimated by phylogenomic analysis. c) Comparison of pairwise distances between leaves in the sequence similarity and gene content trees, for three representative groups (the left-most plot corresponds to the trees in a and b). The red line is the fit of the recombination-driven model of sequence divergence with a long-term delay in the molecular clock equal to *τ*_∞_.

### A recombination barrier model of genome divergence explains the delay in the molecular clock

The observation that early divergence of strains proceeds through a transient stage in which substitutions (effectively) do not accumulate motivated us to study the dynamics of sequence divergence in the presence of homologous recombination. Rather than focusing on the mechanistic details of recombination, we formulated a phenomenological model for the fraction of loci within a genome that become isolated with respect to another genome from the same original population (see Methods). Loci that are not affected by barriers to recombination experience periodical gene conversion that reverts them to the “population average”^21,29^. Our model captures this fact by assuming that recombining loci provide a negligible contribution to the genome-wide sequence divergence. However, as soon as barriers to recombination are established, the affected loci start diverging from the ancestral population at a rate that is proportional to the substitution rate. As a result, the overall sequence divergence at a given time results from the contribution of all loci that are isolated by barriers to recombination, weighted by the time elapsed since the respective locus crossed the barrier.

We show in the Methods that homologous recombination within a population induces a delay in the molecular clock, such that the average divergence of a genome with respect to a member of the same population grows as

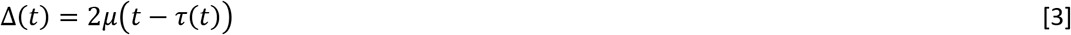

where *t* is the time since the last common ancestor. Mathematically, the delay *τ*(*t*) is a concave and saturating function, the detailed form of which depends on the dynamics of the evolution of recombination barriers (for exact expressions for some simple scenarios, see Supplementary Information). For sufficiently long times, this term reaches a constant value *τ*_∞_, which is the long-term evolutionary delay of the molecular clock induced by homologous recombination. The delay in the molecular clock accounts for the amount of unobserved variation that is erased by gene conversion during the early phases of divergence from the ancestral population.

A notable consequence of the delay in the molecular clock is that the terminal branches of phylogenetic trees inferred from sequence analysis appear shorter than expected given the actual evolutionary times. Specifically, terminal branches are shortened by a distance *μτ*(*t_A_*), whereas internal branches are shortened by a distance *μτ*(*t_A_*) − *μτ*(*t_B_*), where *t_A_* and *t_B_* are consecutive branching times measured from the tips towards the root. Because both *τ*(*t_A_*) and *τ*(*t_B_*) tend to the same value *τ*_∞_ as time passes, the recombination-driven delays cancel out at long evolutionary times and deep internal branches remain approximately unchanged.

To show that recombination-driven delays are the plausible cause for the lack of linearity between the molecular and gene turnover clocks, we used the recombination barrier model to correct the branches of sequence-based trees, accounting for the effects of recombination (see Methods). Then, we sampled subsets of genomes and reassessed the divergence in gene content as a function of the corrected tree distances. Remarkably, correction of the sequence trees led to gene-sharing decay curves that do not depend on the sample size, as predicted by the theory (right panel on Fig. 1c shows a representative case). When extending the same approach to all groups of genomes using taxa-specific delays (see below) we found that the mean separation among the curves obtained for different sample sizes decreased by 30% (permutation test p = 0.012; Supplementary Fig. S2). These results are compatible with the existence of a recombination-driven delay in the molecular clock, which causes a systematic shortening of the terminal branches of phylogenetic trees built from sequence alignments.

### The dynamics of escape from recombination

We leveraged the differences between the substitution and gene turnover clocks to obtain quantitative estimates of the recombination-driven delays in different taxa. Starting from (uncorrected) sequence similarity trees (Fig. 2a) and gene content trees (Fig. 2b), we retrieved all pairwise distances among leaves and plotted the distances from the sequence similarity tree against those from the gene content tree (Fig. 2c). The recombination-driven delays were obtained by fitting the recombination barrier model to such plots. To better understand how barriers to recombination spread along the genome, we evaluated several scenarios for the temporal establishment of such barriers (see Methods). We found that the data in 18 out of the 34 studied groups are best explained by an “autocatalytic” scenario, in which the rate at which barriers spread accelerates as more and more loci become isolated (Supplementary Table S1). The autocatalytic scenario also provides good fits in 13 more groups, although the inclusion of an extra parameter required in this scenario is not statistically justified if the delays are very small. Under the autocatalytic scenario, the fraction of sites susceptible to homologous recombination follows a sigmoidal curve in time, with a relatively sharp transition (with few exceptions) from a state of fully recombining genomes to a state in which all sites freely diverge.

The estimates of the long-term evolutionary delay *τ*_∞_ range from less than 0.01 to more than 0.5 underreported substitutions per site, with broad variation among prokaryotic groups and sometimes even within the same genus (Fig. 3 and Supplementary Table S1). In approximately half of the groups, molecular evolution appears to be strongly delayed, possibly by the pull of recombination, as indicated by the fact that *τ*_∞_ is larger than the depth of the sequence similarity tree. In these cases, there are few pairs of genomes that have reached the regime of free (linear) divergence, and the estimation of the upper 95% confidence bound for the long-term evolutionary delay becomes unfeasible. The 5 groups of Firmicutes included in the analysis (covering bacilli, clostridia and streptococci) belong to this category. In contrast, representatives from the genus *Pseudomonas* are characterized by a linear divergence regime, with little or no signs of recombination-driven delay.

**Figure 3.**
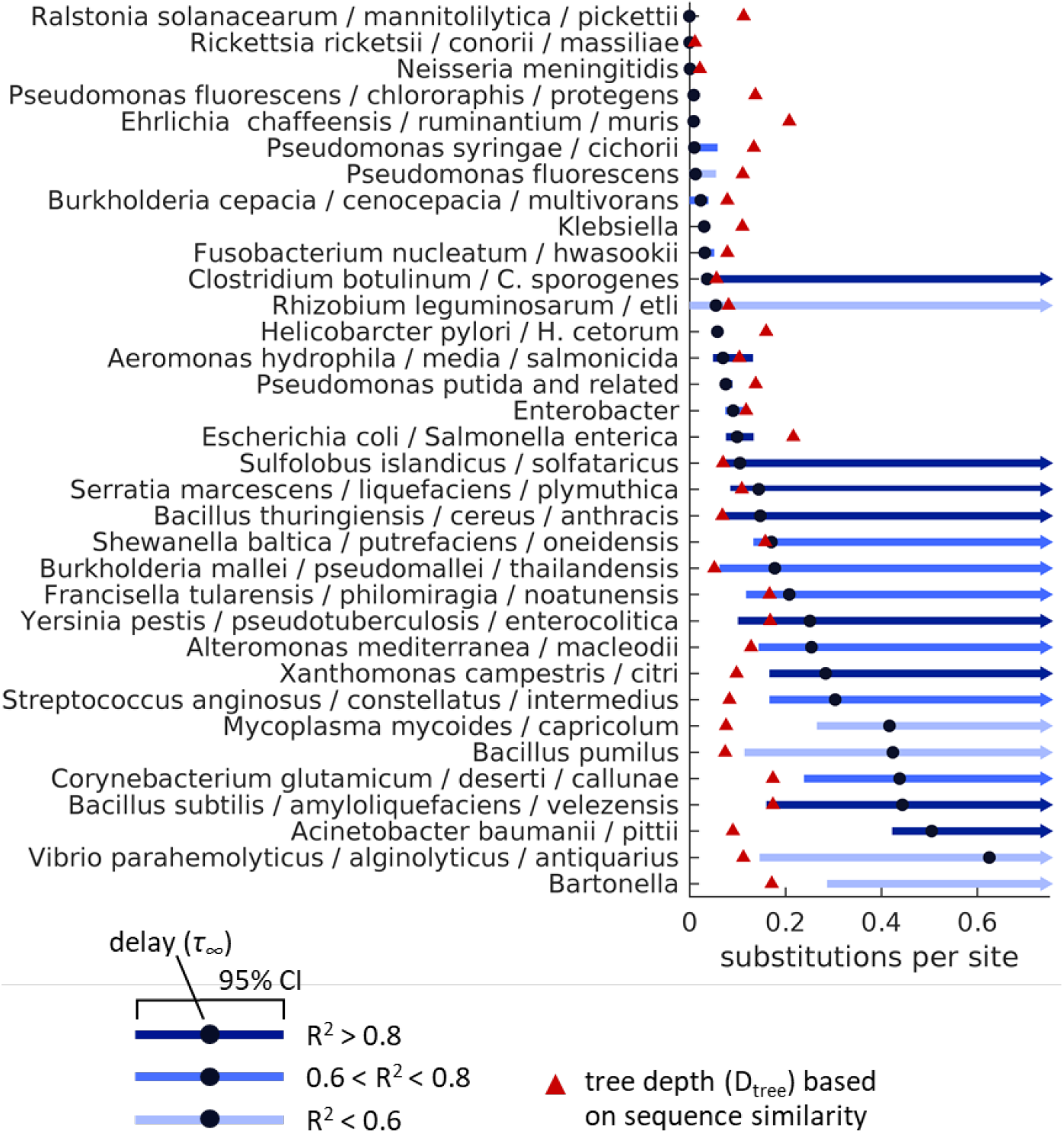
Recombination-driven delay (*τ*_∞_) in the molecular clock in different groups of bacteria. Black circles indicate the best fit of the long-term delay parameter *τ*_∞_ based on the comparison of sequence similarity and gene content trees. Blue lines show the 95% confidence intervals. Red triangles indicate the total depth of the sequence similarity tree. Values of the delay above or very close to the total tree depth imply that most genomes in a group are strongly bound by homologous recombination; an upper 95% confidence bound cannot be calculated in those cases.

The magnitude of the recombination-driven delay is tightly linked to the time frame over which sequence divergence takes place. In taxa with little or no delay, variations in the time at which different genes cross the recombination barrier are negligible; from the perspective of divergence times, all genes in these taxa start diverging roughly at the same time (Fig. 4a, left). Conversely, in taxa with lengthy delays, between-gene variations in divergence times can be comparable to the total evolutionary depth of the taxon (Fig. 4a, right). Accordingly, it can be expected that differences in sequence divergence across genes will be larger in taxa with long recombination-driven delays. To test whether genomic data are compatible with this prediction, we first calculated, for a set of 100 nearly universal gene families, the gene family- and taxa-corrected evolutionary rates (i.e. the gene-specific substitution rates normalized by the gene family- and taxon-averaged substitution rates). It can be shown that the standard deviation of evolutionary rates corrected in this way is equal to the coefficient of variation of the times over which genes have been diverging (see Supplementary Information). As predicted by the model, there is a significant positive correlation between the recombination-driven delays and the variance of evolutionary rates (R = 0.64, p < 0.001). Moreover, when comparing taxa with long and short delays (relative to the evolutionary depth), we found that variance of evolutionary rates is significantly greater in the taxa with long delays (Fig. 4b; p = 0.002, Student’s T-test).

**Figure 4:**
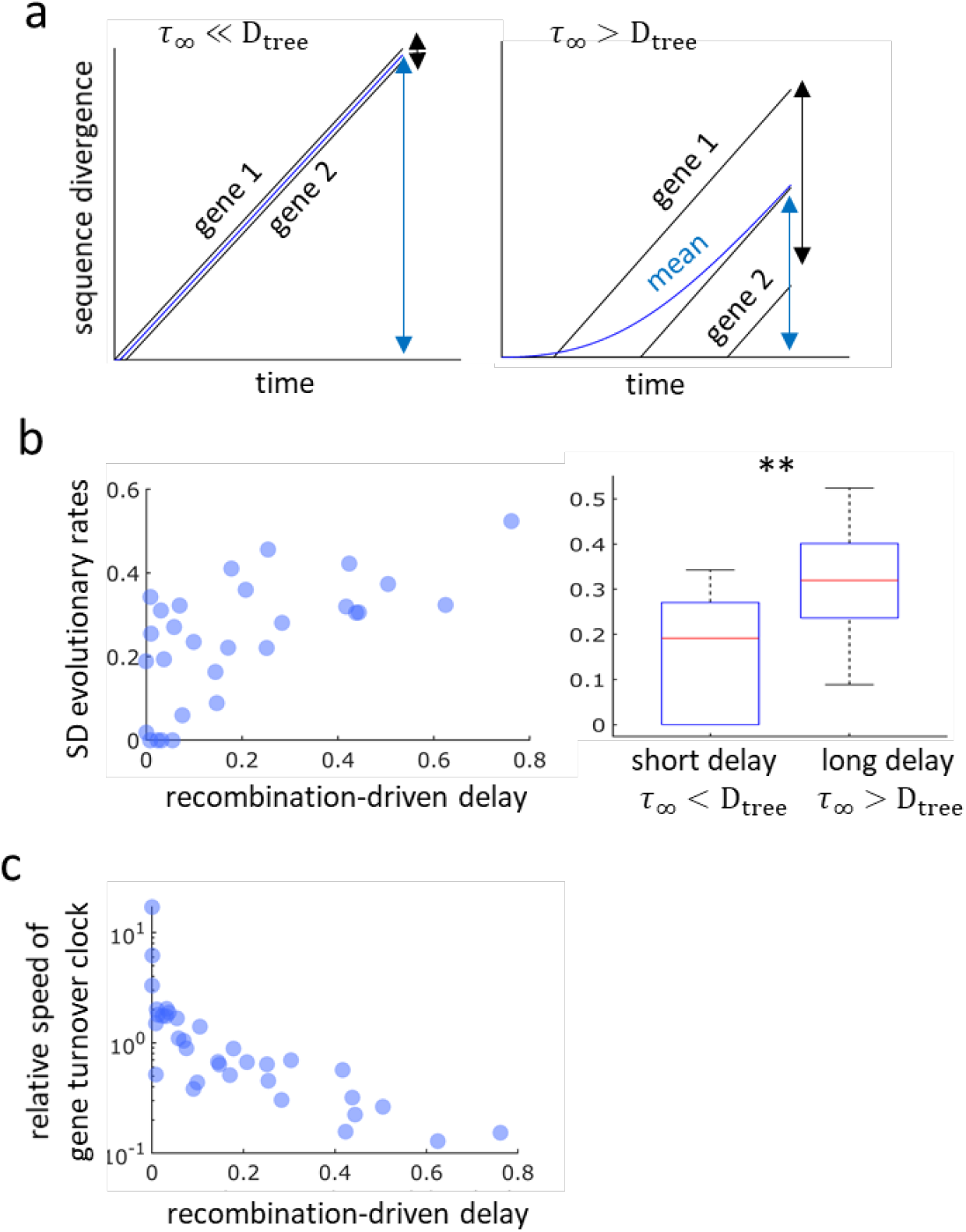
Recombination-driven delay leads to over-dispersion of evolutionary rates within a taxon (a, b) and negatively correlates with the gene turnover rate (c). (a) The divergence of individual genes (black lines) grows linearly after the establishment of barriers to gene conversion, whereas the overall divergence (blue lines) grows non-linearly as the barriers spread across the genome. The mean (blue arrows) and standard deviation (black arrows) of the gene-specific divergences are determined by the values of the recombination-driven delay (τ_∞_) and the tree depth (D_tree_). (b) Standard deviations of the residual evolutionary rates (corrected by gene- and taxon-wise rates) in linearly diverging (τ_∞_ > D_tree_) and strongly delayed taxa (τ_∞_ ≪ D_tree_); Pearson’s correlation coefficient R = 0.64 (p<0.001). ** Statistically significant with p = 0.002 (Student’s T test). (c) Negative association between the recombination-driven delays and the relative rates of gene turnover with respect to substitutions; Spearman’s correlation coefficient rho = −0.86 (p<0.001).

To elucidate potential causes for the large variation in the recombination-driven delays found in the data, we searched for associations with genomic and ecological features, such as genome size, number of mobile genetic elements, gene turnover rate, lifestyle (free-living or host-associated), natural competence for transformation, and effective population size (Supplementary Fig. S3). Among those, we only found a strong negative correlation between the recombination-driven delay and the relative rate of gene turnover with respect to substitutions (Fig. 4c, Spearman’s rho = −0.86, p<0.001), which suggests that fast gene turnover facilitates the spread of barriers to recombination. The statistical power for the analysis of ecological traits (lifestyle and effective population size) was low, and therefore, it cannot be ruled out that these factors contribute to the spread of recombination barriers as well.

## Discussion

Evolution of prokaryotes occurs at two levels: at the sequence level, through substitutions and small indels, and at the genome level, through the transfer and loss of genes and groups of genes^4,30–32^. Whereas evolution at the sequence level is traditionally used to determine phylogenetic relationships among prokaryotes and to assign new genomes to taxonomic groups, it is the gene repertoire (and therefore evolution at the genome level) which determines metabolic capacities, ecological properties and pathogenicity of bacterial strains^33–35^. By studying sequence and gene content divergence in closely related groups of prokaryotes, we found that sequence evolution is often delayed with respect to genome evolution, although the magnitude of the delay broadly varies across bacterial and archaeal lineages. We show that the delay in sequence evolution is likely to result from gene conversion that homogenizes the core genome while, at the same time, accessory genes can be gained and lost. These results are fully compatible with our previous findings indicating that archaeal genomes contain a subset of genes that turn over extremely rapidly, before detectable sequence divergence occurs^26^. The delay on sequence divergence caused by homologous recombination provides a mechanistic explanation for the previously observed “instantaneous” gene turnover in prokaryotes^36^. These results are also compatible with the observation that genes are gained and lost at higher rates on the tips of phylogenetic trees^2^.

Notwithstanding the long-standing debate on the applicability of the species concept in prokaryotes, several recent studies strongly suggest that speciation does occur in bacteria and is a crucial factor shaping the earth microbiome^16,37^. The establishment of barriers to recombination is a pivotal step in the early divergence of closely related prokaryotic strains that eventually leads to speciation^19,23^. In the absence of such barriers, sequence divergence is prevented by the cohesive effect of intra-strain homologous recombination; only after the barriers arise and recombination ceases, substitutions start to accumulate at an appreciable rate. By modeling sequence evolution in the presence of gene conversion, we show here that the temporal dynamics for the spread of barriers to recombination directly affects the molecular clock. Specifically, homologous recombination sets back the molecular clock by an amount of time that, in the long-term, equals the average waiting time for the establishment of recombination barriers. Homologous recombination during early divergence also leads to the compression of the tips of phylogenetic trees. Our model provides a framework for correcting such trees, to make them consistent with the dynamics of gene turnover.

The causes that underlie the spread of barriers to recombination are complex and likely involve genomic and ecological factors. Theoretical models show that barriers can simply emerge if mutations generate diversity faster than recombination erodes it. However, it is a matter of debate whether mutation and recombination alone can explain the formation of non-recombining species in nature^20,21^. Our finding that recombination-driven delays negatively correlate with the rate of gene turnover supports the hypothesis that gene gain and loss facilitates the establishment of barriers to homologous recombination, by promoting niche differentiation^18^ and/or by interfering with gene conversion at flanking loci^22–24^. Moreover, the autocatalytic (or sigmoidal) dynamics that best describes the fraction of recombination-free loci in our model is consistent with the previous findings indicating that barriers to recombination are initiated by the acquisition of lineage-specific genes and subsequently spread from the vicinity of those genes towards the rest of the genome^8^.

Recombination-driven delays are indicative of the time frame over which different genes start to diverge during the split of prokaryotic lineages. Lineages with short delays are characterized by low variance in the gene-specific (gene family- and taxon-corrected) evolutionary rates, which implies that most of their genes began diverging over a brief period of time. In contrast, the higher variances observed in lineages with long delays imply that, in those lineages, divergence of genes occurred in an asynchronous way and spanned longer periods of time. To illustrate this point, it has been estimated that the split between Escherichia and Salmonella took place around 140 million years ago and spanned over 70 million years^8^, which is in close agreement with our finding that the delay in the Escherichia/Salmonella group is approximately half of its evolutionary depth. The high variability of the sequence divergence delays across taxa implies that the time required for lineages to split and form new species is strongly taxon-dependent. For taxa with long delays, it appears natural to think of speciation events being driven by the emergence of strong recombination barriers, whereas taxa with short delays would follow a more continuous speciation dynamics, with new species forming as the genomes gradually diverge^20,21^. Although our analysis focused on prokaryotic genomes, the conclusions appear to be general and can be extended to the evolution of other genomes that are substantially affected by horizontal gene transfer and homologous recombination, for example, viruses.

## Methods

### Recombination-driven delay in sequence divergence

We adopted a phenomenological approach to modeling the effect of intra-population homologous recombination on the genetic diversity of a population. Under this approach, each genomic region is either subject to periodical recombination with other members of the population or has reached a point where homologous recombination is not possible anymore. In the latter case, the respective genomic region is assumed to have crossed (diverged beyond) the recombination barrier. There was no attempt to explicitly model the molecular mechanisms that underlie barriers to recombination, and therefore, the present model is, in principle, applicable to scenarios in which ecological factors, rather than accumulated sequence divergence, drive the isolation of evolving populations. Instead, given a pair of genomes, we used a general function *f*(*t*) to describe the fraction of each genome that is subject to recombination. The probability that a region of the genome crosses the recombination barrier exactly at time *t* (where the time starts counting at the last common ancestor of the two genomes) is

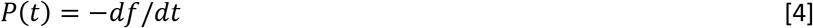

Considering that only the regions that have crossed the recombination barrier make a long-term contribution to sequence divergence, and that the number of substitutions in those regions is proportional to the time elapsed since the recombination barrier was crossed, the overall sequence divergence between a pair of genomes becomes

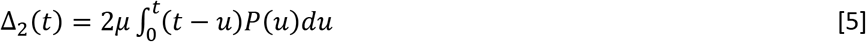

where *μ* is the average substitution rate. Integration by parts leads to the final result

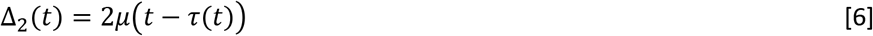

where 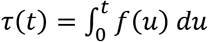 represents a recombination-driven delay in the molecular clock.

To contrast the model with genomic data, we explored three more specific scenarios by assigning explicit functional forms to the rate at which regions of the genome cross the recombination barrier. Thus, we considered a power law time-dependency of the rate *R*(*t*) = *λt^γ^* (which includes the case of a constant rate); a linear time-dependency plus a constant term *R*(*t*) = *λ*_0_ + *λ*_1_*t*; and an “autocatalytic” scenario *R*(*f*) = *λ*_0_ − *λ*_1_*f* in which the rate increases as more and more regions of the genome cross the recombination barrier. Given a functional expression for *R*(*t, f*), the fraction of the genome subject to recombination is obtained by solving the differential equation *df* /*dt* = *R*(*f, t*)*f* (see Supplementary Information for further details).

### Model of genome content divergence

The number of genes, *x*, in a prokaryotic genome was modeled as a stochastic birth-death process, with genome-wide gain rate *P*^+^ and loss rate *P*^− 25^. Under this model, the number of genes in a genome is described by the differential equation

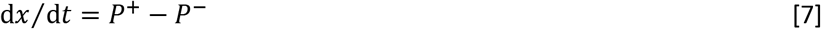

To facilitate comparison of gene content across genomes, each genome was represented by a vector ***X*** with elements that assume values of 1 or 0. Each entry represents a gene, *i.e*. an ATGC-COG, where 1 or 0 indicate the presence or absence, respectively, of that ATGC-COG in the genome. Genome size *x* is then given by the sum of all elements in ***X***.

The number of shared genes in a pair of genomes, or pairwise intersection, *I*_2_ is defined as

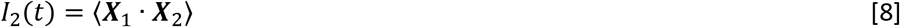

where ***X***_1_ and *X*_2_ are vectors that represent the two genomes, the angled brackets indicate averaging over multiple realizations of the stochastic process, and the dot operation stands for a scalar product. The dynamics of pairwise intersections is given by

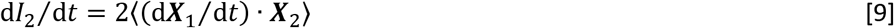

where we used the fact that both averages are equal 〈(d***X***_1_/d*t*) · ***X***_2_〉 = 〈***X***_1_ · (d***X***_2_/d*t*)〉. Assuming that genes are acquired from an (effectively) infinite gene pool ^38^, we have

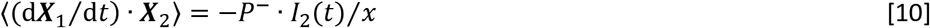

and substituting this relation into the differential equation for pairwise intersections of Eq.(9), we obtain

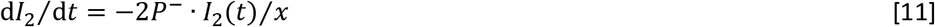

The solution of this equation shows that pairwise genome intersections decay exponentially as

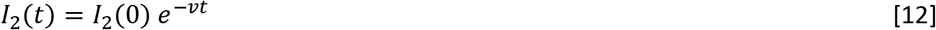

with decay constant *v* = 2P^−^ /*x*. Assuming a molecular clock, the time *t* can be translated into tree pairwise distance as *D*_2_ = 2*t*/*t*_0_ and the pairwise similarity decays exponentially with the tree distance *D*_2_ as

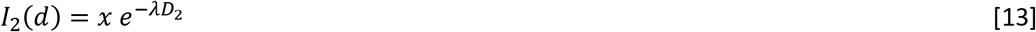

with decay constant *λ*= *t*_0_ P^−^ /*x*. Note that the ratio *P*^−^ /*x* gives the per-gene loss rate.

The intersection of *k* genomes, *I_k_*, is the number of orthologous genes that are shared by all *k* genomes. It is formally defined as

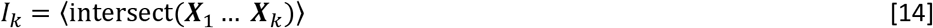

Similar to pairwise genome intersections, the time derivative of *k*-intersections is given by

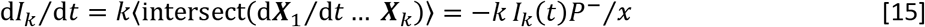

Solving the differential equation, we obtain an exponential decay for the *k*-intersections

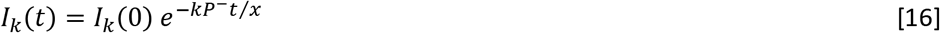

If time is inferred from a tree

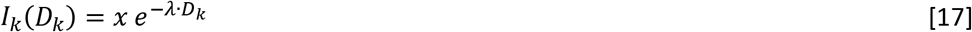

where *λ* is proportional to *P*_−_/*x* and *D_k_* is the sum of branch lengths in the tree that encompasses the *k* genomes (see section 2 of the Supplementary Information for formal derivation). This expression can be extended to genomes composed of fast and slow evolving genes, and becomes

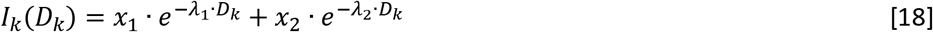

where *x*_1_ and *x*_2_ are the average numbers of genes of each class.

### Genomic data

We used the Alignable Tight Genomic Clusters (ATGC) database^39^ to define groups of closely related bacterial and archaeal genomes. By construction, ATGCs are independent of taxonomic affiliation and meet the objective criteria of high synteny and low divergence (synonymous substitution rate dS<1.5 in protein-coding genes). We selected 36 ATGCs that matched the following criteria: i) maximum pairwise tree distance is at least 0.1 substitutions per site, and ii) the phylogenetic tree contains more than two clades, such that pairwise tree distances are centered around more than two typical values. Two of the 36 genome clusters were identified as outliers and were excluded from the dataset. The 34 genome clusters analyzed in this study are listed in Supplementary Table S1. To facilitate computational analysis, we subsampled large ATGCs to keep at most 20 representative genomes per ATGC. ATGC-specific Clusters of Orthologous Genes (ATGC-COGs) were downloaded from the ATGC database and postprocessed to obtain finer grain gene families by reconstructing approximate phylogenetic trees from original ATGC COG alignments and splitting them into subtrees with minimum paralogy. ATGC-specific phyletic profiles were built by registering, as a binary matrix, the presence or absence of each ATGC-COG in each genome within the ATGC (multiple genes from a single genome that belong to the same ATGC-COG were counted once).

### Tree construction

High-resolution phylogenetic trees based on the concatenated alignments of single-copy core genes were downloaded from the ATGC database^39^. We refer to these trees as sequence similarity-based trees. The phylogenomic reconstruction software Gloome^40^ was used to obtain trees based on the gene content similarity among the members of each ATGC. As the input for Gloome, we used the phyletic profiles for the presence or absence of each ATGC-COG, and the sequence similarity-based trees from the ATGC database; options were set to optimize the tree branch lengths under a genome evolution model with 4 categories of gamma-distributed gain and loss rates. This procedure resulted in 2 trees per ATGC, both with the same topology but with different branch lengths (one based on sequence divergence, the other on gene content divergence). All trees were inspected for extremely long and short branches, and clades responsible for such branches (typically 1 or 2 genomes in 5 out of the 34 ATGCs) were manually removed to avoid possible artifacts in the following steps.

### Tree comparison and model fitting

For each ATGC, we computed all pairwise distances among tree leaves in the sequence similarity- and gene content-based trees. Then, we compared the observed relationship between both sets of distances with the expectations of the recombination barrier model under four scenarios for the recombination barrier crossing rate (constant, linearly increasing with time, linearly increasing plus a constant term, and proportional to the fraction of the genome that has already crossed, which leads to an “autocatalytically” accelerated crossing rate). For each scenario, we fitted the model parameters using non-linear least-squares optimization (implemented in Matlab R2018b), with sequence similarity-based tree distances as independent variable and gene content-tree distances as dependent variable. The choice of sequence similarity-based tree distances as the independent variable was motivated by the need to fulfil the assumption of homoscedasticity in the non-linear regression model. Additionally, we studied the fit of a heuristic power law model *y* = *bx^α^*. To compare the goodness of fit provided by different models, we calculated the Akaike Information Criterion (AIC) as *AIC* = 2*k* + *n*(ln(2*π RSS*/*n*) + 1), where *k* is the number of parameters, *n* is the number of observations, and *RSS* is the residual sum of squares^41^. The 95% confidence interval for the delay parameter *τ*_∞_ was obtained by finding the values of *τ*_∞_ such that the residual sum of squares becomes *RSS* = *RSS\** (1 + 1.96^2^/(*n* − 1)), where *RSS\** is the residual sum of squares produced by the best fit of the model^42^.

### Correction of tree branch lengths

To account for unobserved variation, tree branch lengths were corrected by using the autocatalytic barrier spread model with the parameters inferred in the previous step. In that model, the observable divergence increases with time according to the function

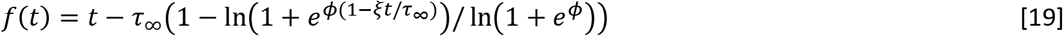

with

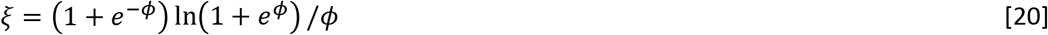

In the simplest case of an ultrametric tree, the corrected height (the distance from the tip) of a node *i*, 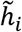, can be calculated by applying the inverse function to the original height *h_i_*, that is

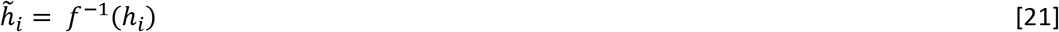

Branch lengths would then be obtained by subtracting the depths of the parent and child nodes. Extending this idea to non-ultrametric trees, we first defined the parental height of a node, *H_i_*, as the distance between the node’s parent and the tip, that is, *H_i_* = *h_i_* + *b_i_*, where *b_i_* is the length of the branch that connects node *i* to its parent node. Parental heights were calculated as weighted averages from the tips to the root, such that *H_i_* = *b_i_* for the leaves and

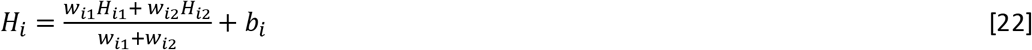

for internal nodes. Subindices *i*1 and *i*2 refer to the child nodes of node *i*; weights were computed as *w_i_* = *b_i_* for leaves and *w_i_* = *w*_*i*1_ + *w*_*i*2_ + *b_i_* for internal nodes (thus, the weight of a node is equal to the total length of the subtree that contains that node as its root plus the length of the branch that connects it to its parent node). Next, corrected values of the parental heights were obtained by computing, numerically (MATLAB R2018b), the inverse function 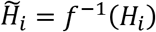. The corrected branch lengths for the tips are simply 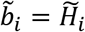. Finally, we proceeded from tips to root and obtained the corrected branch lengths associated with internal nodes as

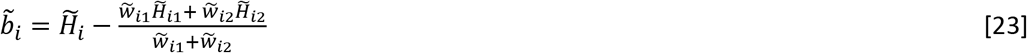

where the new weights 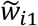 and 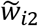 were recalculated at each step using the corrected branch lengths.

### Analysis of gene content

For a set of *k* genomes that belong to the same ATGC, the gene content overlap was calculated as the number of ATGC-COGs shared by all the genomes divided by the mean number of ATGC-COGs per genome. The total sequence divergence of a set of *k* genomes was calculated as the sum of all branch lengths in the sequence similarity subtree that results from selecting the corresponding leaves in the whole-ATGC tree. Curves for the temporal decay of the fraction of shared genes were obtained by plotting the gene content overlap against the total sequence divergence for all possible combinations of *k* genomes within the ATGC. Smooth curves were obtained by fitting a cubic spline model with 5 knots (placed in both extremes and in the 25-, 50-, and 75-percentiles of the data x-values) using the SLM tool (D’Errico, August 10, 2017 (http://www.mathworks.com/matlabcentral/fileexchange/24443)) in MATLAB R2018b. The mean separation among the curves obtained for different values of *k* was calculated as

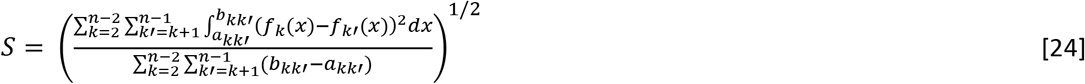

where *n* is the number of genomes in the ATGC, *f_k_* and *f_k′_* are the curves that result from fitting the spline model to sets of *k* and *k′* genomes, respectively, and *a_kk′_* and *b_kk′_* are the bounds of the x-axis interval in which both *f_k_* and *f_k′_* are defined^43^. A value of *S* = 0 corresponds to the exact coincidence of the curves for all values of *k*, which is expected in the absence of evolutionary delays. To assess whether correction of branch lengths for unobserved variation reduces the separation among curves, we calculated the relative change in separation as (*S_original_* − *S_corrected_*)/max(*S_original_, S_corrected_*), which takes the value of 1 when correction leads to complete collapse, positive values ≤1 when correction reduces separation, and negative values ≥-1 when correction increases separation among curves. The statistical significance of the relative change in separation was assessed using a permutation test that involved calculating the median relative change across ATGCs and comparing it with 10^6^ randomized datasets in which the “original” and “corrected” labels were randomly reassigned in each ATGC.

### Quantification of evolutionary rates

For the analysis of evolutionary rates, we focused on a previously published list of 100 nearly universal gene families^44^, defined as clusters of orthologous genes or COGs^45^. To minimize possible confounding effects due to paralogy, we identified all the ATGC-COGs that match any of the universal COGs and restricted the analysis to those COGs that are represented by a single ATGC-COG in at least 30 of the 34 analyzed ATGCs. Multiple sequence alignments for the selected ATGC-COGs were downloaded from the ATGC database and processed to extract all pairwise distances (Nei-Tamura method). Only index orthologs from the ATGC database, i.e. a single sequence per ATGC-COG per genome, were included. For each ATGC-COG, pairwise distances between sequences were plotted against pairwise distances in the phylogenetic tree, and a linear regression model with zero intercept was applied to obtain the relative evolutionary rate of the ATGC-COG with respect to the ATGC average. To minimize the impact of rare instances of gene replacement, which manifest as a non-linear relationship between sequence and tree pairwise distances, we discarded the ATGC-COGs with the fit to the regression model *R*^2^ < 0.9. To account for COG-specific evolutionary rates, the relative evolutionary rate of each ATGC-COG was divided by the mean of all ATGC-COGs that match the same COG. The result is the ATGC-COG residual evolutionary rate, that is, the ATGC-COG evolutionary rate corrected by COG- and ATGC-wise averages. For each ATGC, the dispersion of evolutionary rates was quantified as the standard deviation of the residual evolutionary rates of its ATGC-COGs.

## Author contributions

JI and IS performed research; JI, YIW, EVK and IS analyzed the data; JI, EVK and IS wrote the manuscript that was edited and approved by all authors.

## Acknowledgements

The authors thank Koonin group members fir helpful discussions. The authors’ research is supported by intramural program funds of the National Institutes of Health.

