## Supporting information for "Homologous recombination substantially delays sequence but not gene content divergence of prokaryotic populations"

**2 Genome content divergence** **5**

**3 Delay in sequence divergence driven by homologous recombination** **6**

**4 Analytical study of simple cases** **7**

**5 Dispersion of evolutionary rates** **8**

### 1 Supplementary Figures

#### 1.1 Supplementary figure S1

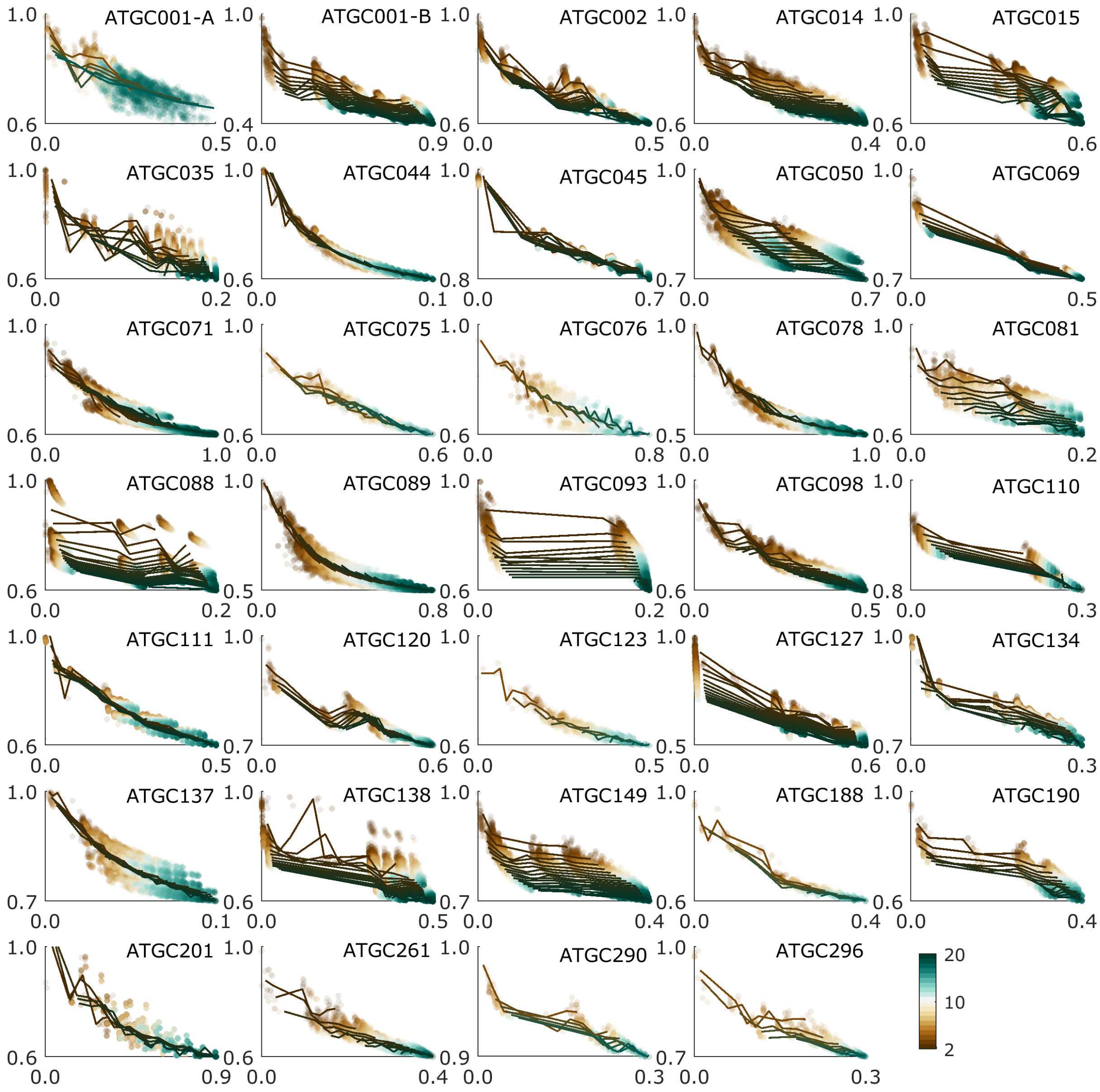

**Figure S1.** Fraction of shared genes as a function of the total tree distance, based on traditional sequence similarity trees. Colors indicate the number of genomes sampled in each case.

#### 1.2 Supplementary figure S2

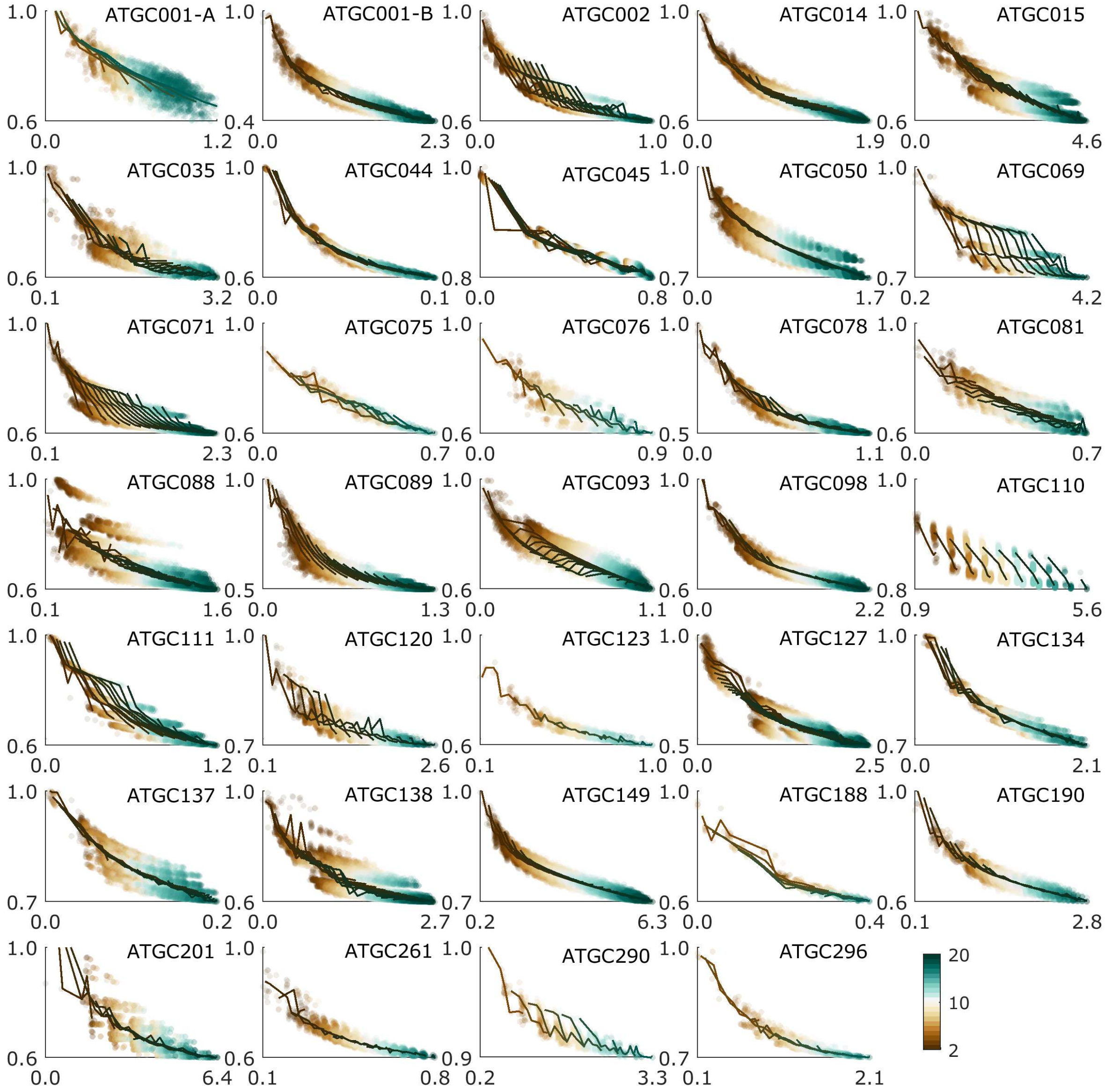

**Figure S2.** Fraction of shared genes as a function of the total tree distance, based on phylogenetic trees whose branch lengths have been corrected to account for the recombination-driven delay in the molecular clock. The corrected branch lengths were obtained by applying the recombination barrier model with the ATGC-specific parameters from Supplementary Table S1 (see Methods). Colors indicate the number of genomes sampled in each case.

#### 1.3 Supplementary figure S3

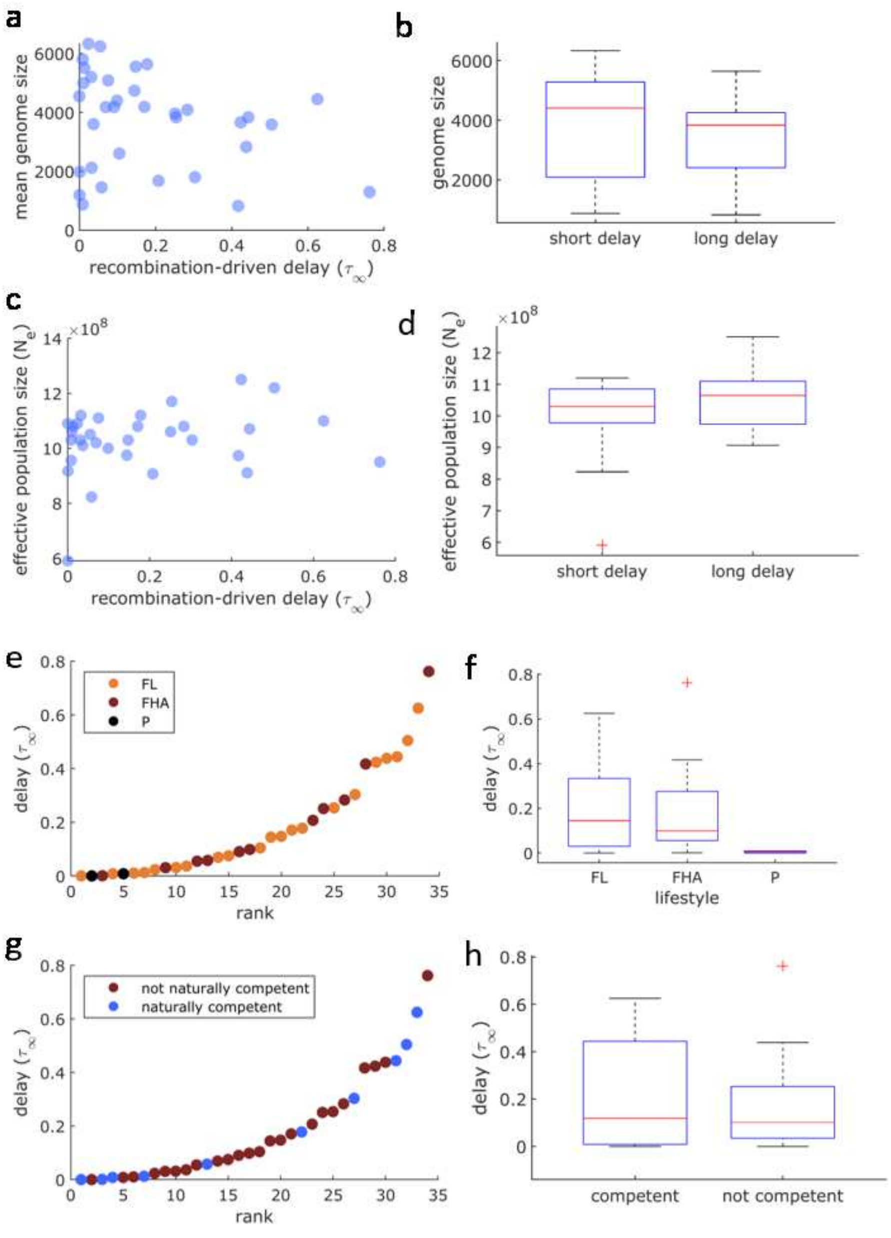

**Figure S3.** Genomic and ecological variables that do not show significant statistical association with the recombination-driven delay ($\tau_\infty$). Each data point corresponds to one ATGC. (a,b) Mean genome size, measured as number of genes; (c,d) effective population size, calculated as described in [2]; (e,f) life style, classified as free-living (FL), facultative host-associated (FHA), and obligate parasite (P); (g,h) natural competence for transformation. Short and long delays refer to the relative magnitude of the delay with respect with the total tree depth.

#### 2 Genome content divergence

The number of genes $x$ evolves under stochastic dynamics that is given by the equation

$$\dot{x} = P_{+} - P_{-} \tag{1}$$

where $P_{+}$ and $P_{-}$ denote gain and loss rates of one gene, respectively. The intersection of $k$ genomes $I_k$ is given by

$$I_k = \langle \text{Intersect}\,(X_1 ... X_k) \rangle \tag{2}$$

where $X_i$ is an array represntaion of a genome that assumes values of 0 or 1 in each entry, indicating presence or absence of genes in each genome. Accordingly, the dynamics of the intersection is given bye

$$\dot{I}_k = k \langle \text{Intersect}(\dot{X}_1 ... X_k) \rangle \tag{3}$$

where the angled brakets indicate averaging on many realizations, and we used the fact that

$$\langle \text{Intersect}(...\dot{X}_n, X_m ...) \rangle = \langle \text{Intersect}(...X_n, \dot{X}_m ...) \rangle \tag{4}$$

Assuming an infinite gene pool, intersections evlove due to loss events, where every time a gene is lost there is a probablity that the lost gene belongs to the intersection, such that

$$\langle \text{Intersect}(\dot{X}_1 ... X_k) \rangle = -P_{-} I_k / x \tag{5}$$

Substituting Eq.(5) into Eq.(3), we get

$$\dot{I}_k = -k P_{-} I_k / x \tag{6}$$

For constant gain and loss rates, the solution of the above equation is given by

$$I_k(t) = I_k(0) \cdot \left[ 1 + \frac{P_{+} - P_{-}}{x} \cdot t \right]^{-\frac{k P_{-}}{P_{+} - P_{-}}} \tag{7}$$

For the genome size equilibrium limit, $(P_{+} - P_{-}) \to 0$ the intersection decays exponentially

$$I_k(t) = I_k(0) \cdot e^{-k P_{-}/x \cdot t} \tag{8}$$

For two genomes the intersecion at $t = 0$ is given by the mean genome size $x$ and we have

$$I_2(t) = x \cdot e^{-\nu \cdot T_2} \tag{9}$$

where the decay constant is given by $\nu = 2P_{-}/x$ and $T_2$ is the time since the latest common ancester of the two intersecting genomes. For three genomes the general result of Eq.(8) takes the from

$$I_3(t) = I_3(0) \cdot e^{-3P_{-}/x \cdot T_3} \tag{10}$$

where $T_3$ measures the time of divergance of three genomes (see Fig.S4). Accordingly, $I_3(0)$ gives the number of commons genes in two genomes right beofre the occurance of a speciasion event that resulted in three intersecting genmoes

$$I_3(0) = x \cdot e^{-2P_{-}/x \cdot T_2} \tag{11}$$

Substituting Eq.(11) into Eq.(10) we get

$$I_3(t) = x \cdot e^{-P_{-}/x \cdot \tau_3} \tag{12}$$

where $\tau_3 = 2T_2 + 3T_3$ is the total evolution time of the three diverging genomes. Assuming a clock, that is, that tree branch lengths are proportional to the time, Eq.(12) can be written in terms of tree distances

$$I_3(t) = x \cdot e^{-\lambda D_3} \tag{13}$$

where $\lambda \propto P_{-}/x$ and $D_3 = \sum d_i$ is the sume of all branch lengths in the tree spanned by the three intersecting genomes. the above result for three intersecting genomes can be generalized to $k$ genomes, leading to the expression

$$I_k(t) = x \cdot e^{-\lambda D_k} \tag{14}$$

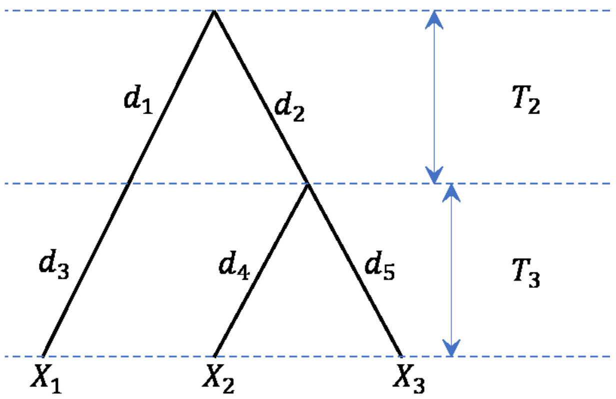

**Figure S4.** Phylogenetic tree depicting divergence of three genomes, noted $X_1$, $X_2$ and $X_3$. Branch lengths are denomted by $d_i$, and times of divergence are indicated. Initially two genomes are diverging for time $T_2$. Following a speciasion event at $T_2$, three genomes are diverging for time $T_3$.

### 3 Delay in sequence divergence driven by homologous recombination

Previous modeling and computational works have shown that mutation accumulation in the presence of homologous recombination is characterized by the existence of "recombination barriers", such that once a genomic region has diverged beyond the recombination barrier, it is unlikely to experience further homologous recombination with the ancestral population. The escape from the recombination barrier is a stochastic process that depends on molecular and ecological details such as the specific mechanism of recombination, the dependence of the recombination efficiency on sequence conservation, or the effective population size. Here we take a phenomenological approach to the dynamics of sequence divergence and define the fraction of the genome that is subject to recombination as $f$. As regions of the genome escape the recombination barrier, the fraction subject to homologous recombination is described by

$$\frac{\mathrm{d}f}{\mathrm{d}t} = -Rf \tag{15}$$

where $R$ is the rate at which regions of the genome escape the recombination rate. $R$ can be a constant (as suggested by [1]) or any arbitrary function of $t$ and $f$. The probability that a region of the genome escapes exactly at time $t$ is

$$P(t) = -\frac{\mathrm{d}f}{\mathrm{d}t} \tag{16}$$

Despite substitutions can occur anywhere in the genome, regions subject to homologous recombination periodically revert to the original sequence and therefore their contribution to overall sequence divergence is minor. Instead we focus on the accumulation of substitutions in regions that escape the recombination barrier, which occurs at a rate $\mu$. For a region that escaped the recombination barrier at time $u$, its contribution to the overall sequence divergence is equal to $\mu(t-u)$. Integrating over all possible escape times we obtain the following expression for the genome-level sequence divergence:

$$\frac{\Delta(t)}{\mu} = \int_0^t (t-u)\,P(u)\,\mathrm{d}u \tag{17}$$

Integration by parts leads to

$$\frac{\Delta(t)}{\mu} = -tF(0) + \int_0^t F(u)\,\mathrm{d}u \tag{18}$$

where $F(u) = \int_0^u P(w)\mathrm{d}w$. It follows from eqn. 16 that $F(u) = -f(u)$. Moreover, assuming an initial state in which the whole genome is subject to homologous recombination, it results that $f(0) = 1$ and the overall sequence divergence becomes

$$\Delta(t) = \mu \left( t - \int_0^t f(u)\mathrm{d}u \right) \tag{19}$$

The integral term defines a delay in the molecular clock driven by the presence of homologous recombination.

### 4 Analytical study of simple cases

In general, the delay term in eqn. 19 depends on the dynamics of escape from the recombination barrier, which itself depends on molecular and ecological factors such as the recombination mechanisms and the effective population size. From a purely phenomenological perspective, exact expressions for the delay term can be obtained by considering possible functional dependences of the escape rate with the time since the last common ancestor and with the fraction of escaped regions.

#### 4.1 Case $R = \lambda t^\gamma$

Here we consider the general case in which the escape rate is proportional to an arbitrary power of the time variable. This includes the particular case of a constant escape rate, as well as cases were the escape rate accelerates or decelerates with time. The differential equation for the fraction of the genome subject to homologous recombination is

$$\frac{\mathrm{d}f}{f} = -\lambda t^\gamma \, \mathrm{d}t \tag{20}$$

with the boundary condition $f(0) = 1$. The boundary condition can only be satisfied if $\gamma > 1$, in which case we get

$$f(t) = \exp\left( -\frac{\lambda \, t^{\gamma+1}}{\gamma + 1} \right) \tag{21}$$

The delay term in eqn. 19 is obtained by integrating the function $f$. Simple analytical expressions are obtained in the special cases $\gamma = 0$ (constant escape rate) and $\gamma = 1$ (linear growth of the escape rate with time).

$$\gamma = 0 \quad \Rightarrow \quad \frac{\Delta(t)}{\mu} = t - \tau\,(1 - e^{-t/\tau}) \quad \text{with} \quad \tau = 1/\lambda \tag{22}$$

$$\gamma = 1 \quad \Rightarrow \quad \frac{\Delta(t)}{\mu} = t - \tau \ \mathrm{erf}\left( \frac{\sqrt{\pi}}{2} \frac{t}{\tau} \right) \quad \text{with} \quad \tau = \sqrt{\frac{\pi}{2\lambda}} \tag{23}$$

where erf is the error function. In these expressions, $\tau$ is the long-term delay in the molecular clock induced by homologous recombination.

#### 4.2 Case $R = \lambda_0 + \lambda_1 t$

When the escape rate consists of a constant term plus a linear term in the time variable, the fraction of the genome susceptible to homologous recombination follows the differential equation

$$\frac{\mathrm{d}f}{f} = -(\lambda_0 + \lambda_1 t)\, \mathrm{d}t \tag{24}$$

with solution

$$f(t) = \exp\left(-t\left(\lambda_0 + \frac{\lambda_1}{2}\,t\right)\right) \tag{25}$$

Substituting eqn. 25 into eqn. 19, the overall sequence divergence becomes

$$\begin{aligned}\frac{\Delta(t)}{\mu} &= t - \int_0^t e^{-u\,\left(\lambda_0 + \frac{\lambda_1}{2}\,u\right)}\,\mathrm{d}u &(26)\\ &= t - \tau_1\, e^{\frac{1}{\pi}\left(\frac{\tau_1}{\tau_0}\right)^2}\left[\mathrm{erf}\left(\frac{1}{\sqrt{\pi}}\frac{\tau_1}{\tau_0} + \frac{\sqrt{\pi}}{2}\frac{t}{\tau_1}\right) - \mathrm{erf}\left(\frac{1}{\sqrt{\pi}}\frac{\tau_1}{\tau_0}\right)\right] &(27)\end{aligned}$$

where $\tau_0 = \frac{1}{\lambda_0}$ and $\tau_1 = \sqrt{\frac{\pi}{2\lambda_1}}$. In the limit $t \to \infty$, the long-term delay in the molecular clock becomes

$$\tau = \tau_1\, e^{\frac{1}{\pi}\left(\frac{\tau_1}{\tau_0}\right)^2}\left[1 - \mathrm{erf}\left(\frac{1}{\sqrt{\pi}}\frac{\tau_1}{\tau_0}\right)\right] \tag{28}$$

##### 4.3 Case $R = \lambda_0 - \lambda_1 f$

This case covers the more realistic scenario in which the escape rate depends on the fraction of the genome that has already crossed the recombination barrier. Intuitively, the probability that a region of interest is surrounded by well conserved regions that can act as seeds for homologous recombination will depend on how many of such well conserved regions still exist in the genome. As a result, the rate at which genomic regions escape the recombination barrier will accelerate as the fraction of regions that are still subject to recombination decreases. A simple model for this scenario is represented by the following differential equation:

$$\frac{\mathrm{d}f}{\mathrm{d}t} = -\lambda_0 f + \lambda_1 f^2 \tag{29}$$

For consistency with the biological interpretation, we impose the constraint $\lambda_0 > \lambda_1 > 0$. The solution of this equation with the boundary condition $f(0) = 1$ is

$$f(t) = \frac{1 + e^{-\phi}}{1 + e^{-\phi(1 - t/\tau)}} \tag{30}$$

where $\phi = \ln\left(\frac{\lambda_1}{\lambda_0 - \lambda_1}\right)$ and $\tau = \phi/\lambda_0$. The expression in eqn. 30 describes a sigmoidal dynamics with a transition from $f = 1$ (homologous recombination affecting the entire genome) to $f = 0$ (total escape from the pull of recombination) around a characteristic time $\tau$. Parameter $\phi$ describes the steepness of the transition, with greater values of $\phi$ implying a faster transition. The substitution of eqn. 30 into eqn. 19 provides the following expression for the overall sequence divergence:

$$\frac{\Delta(t)}{\mu} = t - \tau\,\frac{1 + e^{-\phi}}{\phi}\,\ln\left(\frac{1 + e^{\phi}}{1 + e^{\phi(1 - t/\tau)}}\right) \tag{31}$$

In this case, the long-term delay in the molecular clock becomes $\tau\,(1 + e^{-\phi})\,\ln(1 + e^{\phi})/\phi = \ln\left(\frac{\lambda_0}{\lambda_0 - \lambda_1}\right)/\lambda_1$.

#### 5 Dispersion of evolutionary rates

In section 3 we focused on the genome-level sequence divergence, which results from different regions escaping the recombination barrier at different times. In this section, we will study the rate at which individual regions accumulate mutations. Let $\Pr(D|t)$ be the probability that the sequence divergence in the region of interest is equal to $D$ at time $t$. Assuming that the main source of stochasticity in the divergence process is the time required to escape the recombination barrier, we can write

$$\Pr(D|t) = \begin{cases} 1 - \int_0^t P(u)\mathrm{d}u & \text{if } r = 0 \\ P\left(t - D/\mu\right)/\mu & \text{if } 0 < D \leq \mu t \\ 0 & \text{if } D > \mu t \end{cases} \tag{32}$$

where $P(u)$ is the probability that the region escapes the recombination barrier exactly at time $u$.

Let us now introduce the relative evolutionary rate, $r$, as the ratio between the divergence of a region and the genome-level divergence at a given time:

$$r = \frac{D}{\langle D|t\rangle_G} \tag{33}$$

The angle brackets $\langle\cdot\rangle_G$ indicate the average within a genome. Such average affects both the escape times and the substitution rates, because substitution rates can vary across regions. Assuming that the escape times are independent of the substitution rates and applying eqn. 17, it results that

$$\langle D|t\rangle_G = \langle\mu\rangle_G \int_0^t (t-u)P(u)\mathrm{d}u \tag{34}$$

In closely related genomes with a high degree of synteny, the variability in substitution rates across regions can be accounted for by normalizing by the average evolutionary rates of the genes that occupy each region. To that purpose, we define the residual evolutionary rate as:

$$\delta = \frac{r}{\langle r|t\rangle_L} = \frac{D}{\langle D|t\rangle_G \langle r|t\rangle_L} \tag{35}$$

where $\langle\cdot\rangle_L$ denotes the average for a specific locus across closely related genomes. The value of $\langle r|t\rangle_L$ can be calculated by taking the average in eqn. 33 and under a constant value of $\mu$:

$$\langle r|t\rangle_L = \frac{\langle r|t\rangle_L}{\langle D|t\rangle_G} = \frac{\mu}{\langle\mu\rangle_G} \tag{36}$$

and combining eqns. 35 and 36 we get to the final expression for the residual evolutionary rate:

$$\delta = \frac{D\,\langle\mu\rangle_G}{\mu\,\langle D|t\rangle_G} \tag{37}$$

Let $\Pr(\delta|t)$ be the probability that the residual evolutionary rate of a genomic region is equal to $\delta$ at time $t$. An explicit expression for $\Pr(\delta|t)$ can be obtained by substituting eqn. 37 in eqn. 32:

$$\Pr(\delta|t) = \begin{cases} 1 - \int_0^t P(u)\mathrm{d}u & \text{if } \delta = 0 \\ P\left(t - \frac{\delta\langle D|t\rangle_G}{\langle\mu\rangle_G}\right)\frac{\langle D|t\rangle_G}{\langle\mu\rangle_G} & \text{if } 0 < \delta \leq \frac{t\,\langle\mu\rangle_G}{\langle D|t\rangle_G} \\ 0 & \text{if } \delta > \frac{t\,\langle\mu\rangle_G}{\langle D|t\rangle_G} \end{cases} \tag{38}$$

The genomic variance of $\delta$ at a given time is calculated as $\mathrm{Var}(\delta|t) = \langle(\delta|t)^2\rangle_G - \langle\delta|t\rangle_G^2$. The quadratic mean is equal to

$$\langle(\delta|t)^2\rangle_G = \int_0^{\frac{t\,\langle\mu\rangle_G}{\langle D|t\rangle_G}} \delta^2\, P(\delta|t)\,\mathrm{d}\delta = \frac{\langle\mu\rangle_G^2}{\langle D|t\rangle_G^2}\int_0^t (t-u)^2 P(u)\mathrm{d}u \tag{39}$$

The mean $\langle\delta|t\rangle_G$ is equal to

$$\langle\delta|t\rangle_G = \int_0^{\frac{t\,\langle\mu\rangle_G}{\langle D|t\rangle_G}} \delta\, P(\delta|t)\,\mathrm{d}\delta = \frac{\langle\mu\rangle_G}{\langle D|t\rangle_G}\int_0^t (t-u)P(u)\mathrm{d}u = 1 \tag{40}$$

Substituting eqn. 34 into eqn. 39, the genomic variance ot the residual evolutionary rates becomes:

$$\mathrm{Var}(\delta|t) = \langle(\delta|t)^2\rangle_G - \langle\delta|t\rangle_G^2 = \frac{\int_0^t (t-u)^2 P(u)\mathrm{d}u}{\left(\int_0^t (t-u)P(u)\mathrm{d}u\right)^2} - 1 = \frac{\mathrm{Var}(t_{free}|t)}{\langle t_{free}|t\rangle_G^2} \tag{41}$$

where the new variable $t_{free}$ (the free-evolving time) represents the time elapsed since a region escaped the recombination barrier. Therefore, the standard deviation of the residual evolutionary rates is equal to the coefficient of variation of the free-evolving times, which is small if all the genomic regions escaped the recombination barrier a long time ago, and large if parts of the genome are still under the homogeneizing effects of homologous recombination.
